# Stromal COL12A1 defines a pro-invasive extracellular matrix niche in appendiceal mucinous neoplasms and pseudomyxoma peritonei

**DOI:** 10.64898/2026.10.26.761946

**Authors:** Milly Upton, Raghavendar Nagaraju, Angelica Castaneda, Luca Varinelli, Jordi Martinez-Quintanilla, Manuela Gariboldi, Jorge Barriuso, Bipasha Chakrabarty, Sarah T O Dwyer, David Thornton, Omer Aziz, Patrick Caswell, PMPnet group

## Abstract

Pseudomyxoma peritonei (PMP) is an uncommon peritoneal malignancy, most often arising from appendiceal mucinous neoplasms (AMNs), in which tumour progression is accompanied by extensive mucus accumulation. Although PMP is clinically dominated by peritoneal dissemination, the stromal matrix cues that support invasion and metastatic niche formation remain poorly defined. Here, we profile extracellular matrix niche from PMP patient samples, and show that AMNs and peritoneal metastases (PMs) exhibit marked ECM remodelling, including enrichment of mucins, fibrillar matrix components and stromal regulators of collagen architecture. Three-dimensional matrix models demonstrate that PMP cell invasion is strongly influenced by ECM composition, with collagen/fibronectin-rich environments promoting invasion and selected small leucine-rich proteoglycans (SLRPs) attenuating this response. Collagen organisation analysis reveals increased matrix deposition, density and alignment in metastatic lesions, consistent with development of a remodelled pro-invasive stromal niche. Among altered matrix proteins, COL12A1 is prominently upregulated in both primary and peritoneal disease and is enriched within the stromal compartment. Functional co-culture experiments further show that COL12A1-expressing cancer-associated fibroblasts (CAFs) enhance PMP cell invasion, whereas COL12A1 silencing in CAFs reduces this pro-invasive effect. These findings identify stromal COL12A1 as a key component of the PMP matrix niche and suggest that collagen XII-dependent remodelling of the tumour microenvironment may contribute may support dissemination and establishment of the metastatic disease. Together, our data indicate that human AMN and PMP have a compositionally and architecturally remodelled extracellular matrix, within which stromal COL12A1 supports CAF-driven tumour-cell invasion.

## Introduction

Pseudomyxoma peritonei (PMP) is a condition defined by accumulation of mucus and mucus-producing tumour cells within the peritoneal cavity, with an estimated incidence of 3.2 people per million per annum (1, 2), often arising from low- or high- grade appendiceal mucinous neoplasms (AMN). AMNs are slow growing epithelial neoplasms, but when left untreated can lead to the characteristic features of PMP including solid metastases within the peritoneal cavity following rupture of the appendix (3). The current standard therapy for PMP is complete cytoreductive surgery (CRS) followed by intraoperative hyperthermic intraperitoneal chemotherapy (HIPEC) which can result in a 10-year overall survival of 63 % if successful (4), but if unsuccessful or inoperable due to extensive disease or relapse, treatment options are limited with no effective targeted therapies available. Therefore, enhancing our biological understanding of PMP and gaining insight into the environment that surrounds and supports these neoplasms will create opportunity for improved pre-clinical models and novel therapeutic targets for these patients.

The peritoneal membrane forms a barrier lining the abdominal cavity and consists of a monolayer of mesothelial cells, covered by glycocalyx, upon a thin basement membrane extracellular matrix (ECM), beneath which lies a fibrillar collagen-rich interstitial matrix (5, 6). Peritoneal metastases (PM) are a feature of several cancers, often originating from gastrointestinal or gynaecological malignancies (7–9). AMNs show an ability to leave the primary site and disseminate within the peritoneal cavity, however their metastasis is almost always limited to this anatomical site. An increasing body of evidence points towards the ECM within the tumour microenvironment (TME) as a key mediator of metastatic dissemination, with ECM remodelling central to PM niche development as disseminating cells attach to and invade into the peritoneum (6).

The ECM is a highly complex 3-dimensional (3D) meshwork of proteins, glycoproteins and polysaccharides that support the structural integrity of tissues, cell-matrix communications, immune cell mobility and tissue repair through the formation of a scaffold. The matrisome is comprised of core ECM proteins (collagens, glycoproteins (e.g. laminin, fibronectin) and proteoglycans (e.g. biglycan, lumican)) and ECM-associated proteins (ECM-affiliate proteins (e.g. mucins, C-type lectins), ECM regulators (e.g. matrix metalloproteases, lysyl oxidases) and secreted factors (e.g. Wnt ligands, cytokines)) (10). However, within the TME the matrix is often replaced with an aberrant tumour-specific matrix, primarily deposited via cancer-associated fibroblasts (CAFs). This matrix underpins biophysical and biochemical changes that have direct implications on complex cellular signalling networks (11, 12). Many malignant cells secrete various pro-fibrotic growth factors and signalling molecules such as transforming growth factor-beta (TGF-β) and epidermal growth factor (EGF) resulting in the recruitment and differentiation of stromal cells in the TME (13, 14). This in turn leads to the deposition of an abundant and remodelled matrix with altered properties that facilitates both tumour growth and metastatic dissemination (15).

Fibrillar collagens are a family of triple helical proteins that polymerise to form the major structural component of the ECM, and these fibrillar arrays are further regulated through interactions with fibril-associated collagens with interrupted triple helices (FACIT) collagens (including collagen IX, XII and XVI), which organise matrix and determine tissue biomechanics (16, 17). By binding to collagen I fibrils, secreted FACIT collagens increase collagen bundle width and alignment, remodelling a dense fibril meshwork into highly ordered fibres for cancer cells to invade along (17). Alongside collagens, proteoglycans and glycoproteins decorate the matrix to confer both biophysical and signalling properties (18–20). The interplay between CAFs and collagen deposition, remodelling and post-translational modification has consistently been associated with poor prognosis and treatment resistance in various solid tumours such as ovarian and colorectal cancer (CRC) (21–23), and COL1A1 expression in CRC is significantly associated with metastatic dissemination and increased rate of relapse (24). Moreover, collagen orientation and fibre alignment enhance cell motility, cell spreading and focal adhesions in cancer cells suggesting stromal matrix architecture and not just abundance is crucial for the development of a tumour-associated matrix environment in the peritoneal cavity in ovarian cancer (16, 25). However, the composition and role of matrix in AMN and PM is unknown, and how specific components of the complex matrix niche contribute to dissemination is only beginning to emerge.

By profiling the tumour matrisome through matrix protein enrichment coupled with quantitative mass spectrometry in both the primary AMNs and PMs from patients with PMP, we revealed the tumour matrix niche. The matrix was predominantly a fibrillar collagen rich interstitial matrix, and by reconstructing the matrix *in vitro* we demonstrate that some abundant matrix glycoproteins (fibronectin) had a pro-invasive effect on PMP cells, whereas other abundant proteoglycans (biglycan, lumican) had anti-invasive properties. We also observed a striking increase in the FACIT collagen COL12A1 in both primary AMN and PMs. Recent studies highlight the expression of collagen XII as a poor prognostic biomarker in CRC, and a role for collagen XII deposition by CAFs in the development of breast cancer metastases to the lung through the regulation of collagen type 1 fibril organisation (17, 26). We show that knockdown of COL12A1 in PM-derived CAFs prevents the ability of CAFs to support PMP invasion. These data suggest that the profile of insoluble matrix components in PMP could determine peritoneal dissemination and disease progression.

## Materials and methods

### Study approvals

Ethical approval and written patient consent were obtained before the collection of patient samples and was granted by Manchester Cancer Research Centre (MCRC) Biobank Study (Human Tissue Authority licence number 30004); South Manchester Research Ethics Committee 18/NW/0092. All samples were anonymised.

### Patient samples

9 patients diagnosed with PMP were included within this study and samples were resected from patients undergoing cytoreductive surgery at the Colorectal and Peritoneal Oncology Centre, The Christie NHS Foundation Trust, United Kingdom. Following histological confirmation by a Pathologist specialising in PMP, samples were processed for mass spectrometry using snap frozen tissues including uninvolved regions (uninvolved appendix (n=4) and peritoneum (n=4)) versus diseased regions (AMN (n=3) and PM (n=4)). Furthermore, formalin-fixed paraffin embedded (FFPE) sections were obtained to validate identified proteins. In addition, for isolation of CAFs, samples were generated from PMs (n=2). Anonymised patient clinico-pathological features are listed in Supplementary Table 1.

### Enrichment of matrisome proteins

Samples were enriched for predominantly insoluble matrisome proteins using chemical fractionation and sonication. Tissues were cut into small pieces (roughly 1mm^3^) on ice and washed in ice-cold PBS. Tissue homogenates were incubated under gentle rotation for one hour at 4 °C in a 5:1 (v/w) ratio of Tris-lysis buffer (10 mM Tris pH 8 (Fisher Scientific), 150 mM Sodium Chloride (NaCl) (Sigma-Aldrich), 1 % Triton-X 100 (Sigma-Aldrich), 25 mM Ethylenediaminetetraacetic acid (EDTA) (Fisher Scientific), LC-MS grade water (Thermo-Fisher Scientific), and EDTA-free Roche complete protease inhibitor cocktail (Sigma-Aldrich)) to tissue. The supernatants were collected and stored at - 80°C, which were termed soluble cellular protein fractions. The pellets were incubated under gentle rotation for one hour at 4 °C in a 5:1 (v/w) ratio of alkaline detergent buffer (20 mM Ammonium Hydroxide (NH_4_OH, Sigma-Aldrich), 0.5 % Triton-X 100, and phosphate-buffered saline (PBS) (Thermo Fisher Scientific)) to tissue. Samples were centrifuged for 10 minutes at 14,000 g and the supernatants were collected and stored at -80 °C which contained less concentrated soluble cellular proteins. The remaining pellets were resuspended in 50 μl of 1x TEAB/SDS lysis buffer (2x buffer – 100 mM triethylammonium bicarbonate (TEAB) pH 8.5 (Sigma-Aldrich), 10 % sodium dodecyl sulfate (SDS) (Sigma-Aldrich), and LC-MS grade water) and termed ECM fractions. The ECM fractions were then transferred into Covaris AFA glass tubes and sonicated using a Covaris LE220+ Focused Ultrasonicator (Covaris). Sonication parameters were as follows: set duration = 50 seconds/sample, peak power = 500 W, duty factor = 20 %, 200 cycles/burst, and average power = 200 W. Sonicated ECM fractions were then placed on ice.

### Matrisome protein digestion and peptide desalting

Sonicated ECM fractions were reduced with 5 mM dithiothreitol (DTT) (Fisher Scientific) for 10 minutes at 60 °C and alkylated with 15 mM iodoacetamide (IAM) (Sigma-Aldrich) for 30 minutes in the dark. The residual alkylation reaction was quenched using 5 mM DTT and protein concentration was quantified using a Millipore Direct Detect spectrometer (Millipore). Subsequently, 40 μg of sample was prepared for S-Trap column digestion using 1:10 (v/v) 12 % aqueous phosphoric acid (Acros Organics) and 1x S-Trap binding buffer (6x = 100 mM TEAB pH 7.1 and 90 % methanol (Thermo Fisher Scientific)). Samples were loaded onto S-Trap columns (ProtiFi) and digested in a static thermomixer at 47 °C for 1 hour with 1 μg of sequencing-grade modified trypsin (Promega). Peptides were eluted from the column using 30 % acetonitrile (Thermo Fisher Scientific) and 0.1 % formic acid aqueous solution (Fisher Scientific) and collected by centrifugation at 4000 g for 2 minutes. Samples were then desalted using POROS R3 beads (Life-Technologies) and dried using vacuum centrifugation.

### Mass spectrometry data acquisition

Peptides were analysed using liquid chromatography tandem mass spectrometry (LC-MS/MS) using an Orbitrap Elite mass spectrometer (Thermo Fisher Scientific) in the Biological Mass Spectrometry core facility, University of Manchester. Peptides were automatically selected for fragmentation by data-dependent analysis and analysed in one batch. All samples were analysed using 2 hour runs and LC column was equilibrated with a pooled sample consisting of an equal mix of all samples.

### Mass spectrometry data analysis

Initial analyses of the spectral raw file outputs were normalised and quantified using MaxQuant software (version 2.6.7.0) which employs label-free quantification (LFQ) to normalise peptide abundance against the human UniProt database. Default MaxQuant parameters were used including fixed cysteine carbamidomethylation modifications and variable methionine oxidation, protein N-terminus acetylation and lysine biotinylation modifications. LFQ intensities from MaxQuant analysis were normalised, imputed and analysed using the web-based analysis suite “Manchester Proteome Profiler” (27).

### Matrisome data analysis

Matrisome AnalyzeR is a web application that annotates, classifies and tabulates matrisome molecules according to matrisome divisions and categories (28). LFQ intensities were analysed using this software and categorised into: Core matrisome proteins (glycoproteins, collagens and proteoglycans) and matrisome-associated proteins (ECM-affiliated, ECM regulators, and secreted factors).

### Cell culture

N14A human PMP cell line (Roberts et al. 2015) was maintained in Dulbecco’s Modified Eagle’s Medium (DMEM) with 4500 mg/L glucose (D6171, Sigma-Aldrich), 1x glutamax (Gibco), 25 mM HEPES, 5 ug/ml Insulin, 10 % (v/v) fetal calf serum (FCS) (all Sigma-Aldrich), and 1 % (v/v) Antibiotic-Antimycotic (ThermoFisher Scientific). Cells were incubated at 37 °C in a humidified 5 % (v/v) CO_2_ atmosphere and passaged every 3-5 days.

### Isolation of cancer-associated fibroblasts

Patient-derived CAFs were isolated from peritoneal metastases (n=2) as follows. Tissues were minced into small fragments, and enzymatic digestion was performed using 1 mg/ml Collagenase (Sigma-Aldrich) in DMEM containing 1 % (v/v) Antibiotic-Antimycotic for 30 minutes at 37 °C and pipetted every 5 minutes to dissociate cells. Digestion was inhibited using DMEM containing 10 % (v/v) FCS and 1 % (v/v) Antibiotic-Antimycotic and centrifuged at 1500 rpm for 5 minutes to remove the media. The cell pellet was filtered (100 µm pore size; BD Sciences) and dissociated cells were washed through using fresh DMEM. The remaining unfiltered tissue pieces were cultured in DMEM containing 4500 mg/L glucose, 10 mM L-glutamine, 25 mM HEPES, 5 µg/ml Insulin, 10 % (v/v) FCS, and 1 % (v/v) Antibiotic-Antimycotic. After 5 days, tissue fragments and debris were removed and fresh media added. Fibroblast cultures were expanded, with a proportion cryogenically stored in Bambanker^TM^ freezing media (NIPPON Genetics) and a proportion used for characterisation. The remaining fibroblasts were expanded for further experiments.

### Transient transfection

Two rounds of lipofection (Lipofectamine 2000, Invitrogen) were performed to transiently introduce SiRNA into CAF cells, according to the manufactures’ instructions. The following silencer select SiRNA reagents were used: COL12A1 #1 (4392420, ID: s3348) and #2 (4392420, ID: s3349) (Invitrogen).

### Immunofluorescence characterisation

Cells grown on 0.1 % gelatin-coated (Sigma-Aldrich) glass coverslips were fixed with 4 % (w/v) paraformaldehyde (PFA) (Sigma-Aldrich) for 20 minutes at room temperature. Cell membranes were permeabilised with 0.2 % (v/v) Triton-X 100 in PBS without CaCl_2_ and MgCl_2_ (Sigma-Aldrich) and blocked in 1 % (w/v) heat-denatured bovine serum albumin (BSA) (Sigma-Aldrich) for 1 hour at room temperature. Cells were stained with the following primary antibodies: anti-α-smooth muscle actin (SMA) (mouse monoclonal antibody (mAb), A2547, Sigma-Aldrich), anti-fibroblast activation protein (FAP) (rabbit polyclonal antibody (pAb) ab53066, Abcam), and anti-COL12A1 (Rabbit pAb, ab121304,

Abcam). Following labelling, cells were incubated with secondary antibodies including: phalloidin Alexa Fluor 647 (A22287 Life Technologies), Rabbit Alexa Fluor 488-conjugated, Mouse Alexa Fluor 647-conjugated and Hoechst 33258 (ThermoFisher) for 1 hour at room temperature and mounted with Prolong Gold Anti-Fade Reagent (Invitrogen). Images were acquired using a Leica TCS SP8 AOBS inverted confocal microscope using a 20× or 40× APO objective. The confocal settings were as follows, pinhole (1 airy unit), scan speed (1000Hz unidirectional), and format (1024 x 1024). Images were collected using hybrid detectors with the appropriate detection mirror settings and sequential image collection, capturing z-stacks using LAS X software (Leica). Maximum intensity projections of these 3D stacks are shown in the results.

### 3D spheroid invasion assay

Spheroids were generated using N14A immortalised cells or co-cultures of N14A with CAF cultures in 3:1 ratio using the hanging-drop method. In brief, N14A cells were pre-labelled with Vybrant DiO cell-labelling solution (Invitrogen) and CAFs with Vybrant DiD cell-labelling solution according to manufactures instructions. Fluorescently labelled cells were suspended at a 4:1 ratio of complete DMEM to methocel solution (12 mg/ml methylcellulose (Sigma-Aldrich) diluted in complete DMEM). Droplets of cell suspensions at a density of 7500 cells/drop were added to the under-side of a petri dish lid, inverted onto the base of a petri dish and incubated for 24 hours at 37 °C in a humidified 5 % (v/v) CO_2_ atmosphere. Spheroids were injected into collagen hydrogels generated using 3 mg/ml Collagen Type 1 (Rat tail high concentration) (Corning), 8 mM NaOH, 10x PBS, and DMEM media. Additional gel components were added in various combinations including fibronectin (Sigma-Aldrich), biglycan (R&D Systems), and lumican (R&D Systems). Hydrogels were incubated for 20 minutes at 37 °C in a humidified 5% (v/v) CO_2_ atmosphere to allow the gels to solidify and then cultured in full DMEM media supplemented with human recombinant EGF (Sigma-Aldrich) and recombinant human HGF (Peprotech). Spheroids were incubated at 37 °C to a total incubation time of 24 hours before imaged using a Leica TCS SP8 AOBS upright confocal microscope using a 10x APO objective. The confocal settings were as follows, pinhole (1 airy unit), scan speed (1000 Hz unidirectional), and format (1024 x 1024). Images were collected using hybrid detectors with the appropriate detection mirror settings and sequential image collection, capturing z-stacks using LAS X software (Leica). Maximum intensity projections of these 3D stacks are shown in the results. Percentage area of invasion was measured using Image J and determined by measuring the threshold intensity of the core spheroid area and subtracting the value from the threshold intensity of the total area invaded following 24-hour invasion into the hydrogels. A minimum of 3 spheroids per internal triplicate were analysed in a minimum of 3 repeats.

### Picrosirius red staining, microscopy and TWOMBLI analysis

FFPE sections cut to 5 μm were stained with picrosirius red (0.1 % sirius red F3BA in saturated aqueous picric acid at pH 2) for 1 hour and then cleared in 0.1 % acidified water using the Leica ST5010 Autostainer through the Histology Core Facilities, University of Manchester. Collagen-associated birefringence was visualised and triplicate regions of interest (ROI), that were identified from H&E-stained sections by a pathologist per patient, were imaged using an Olympus BX63 upright microscope with polarising filters using a 20x or 40x objective and white-balanced using a DP8- camera (Olympus) in colour mode through CellSens Dimension v1.16 (Olympus). Using Fiji ImageJ, 3 representative areas measuring 100x100 μm per sample were selected based on their location within the stroma and proximity to tumour rich regions. Collagen matrix patterns were analysed using The Workflow Of Matrix BioLogy Informatics (TWOMBLI) pipeline as described (29).

### Hyperion mass cytometry

FFPE sections cut to 5 μm underwent deparaffinization by submerging in xylene for 2x 10 minutes before passing through graded alcohols into distilled water for 5 minutes. After rehydration, slides were incubated in Tris-EDTA pH 8.5 for 30 minutes at 96 °C. Once cooled, slides were washed in PBS and blocked in 3 % (w/v) BSA for 30 minutes at room temperature. All slides were stained with unconjugated antibodies (anti-COL12A1 (Rabbit pAb, ab121304, Abcam) and Cytokeratin 20 (Mouse mAb, Ks20.8, Invitrogen)) overnight at 4 °C. Slides were washed in PBS and incubated for 1 hour at room temperature with lanthanide-conjugated secondary antibodies including: Alexa Fluor goat anti-rabbit 488 and Alexa Fluor goat anti-mouse 488 (Invitrogen) conjugated by the Flow Cytometry Core Facility, University of Manchester to lanthanide metals 160Gd and 165Ho respectively. Following thorough washing, all slides were incubated with the 1-hour specific antibody mastermix of lanthanide-conjugated primary antibodies diluted in PBS with 0.05 % TWEEN and 3 % BSA followed by the overnight specific antibody mastermix of lanthanide-conjugated primary antibodies (Supplementary Table 5). Slides were washed in PBS with 0.1 % Triton-X 100, nuclear stained with iridium (Intercalator-Ir, Standard BioTools) for 30 minutes at room temperature and briefly washed in ultrapure water before air drying. Triplicate representative ROIs were identified from H&E-stained sections by a pathologist, each measuring 1x1 mm. Imaging mass cytometry (IMC) imaging was acquired using a Hyperion imaging mass cytometer (Standard BioTools) according to manufacturer’s instructions. In brief, the ROIs were laser-ablated and ionised into its constituent ions by passing through a plasma source. Each lanthanide-conjugated antibody was detected using time-of-flight mass spectrometry followed by image reconstruction based of metal abundance at each pixel. ROIs were viewed using MCD Viewer (Standard BioTools) and pre-processed using the publicly available IMC- data preprocessing pipeline (https://github.com/BodenmillerGroup/ImcSegmentationPipeline).

Using Fiji ImageJ, 3 representative areas measuring 100x100 μm per sample were selected based on their location within stroma (SMA+)/ tumour rich (CK20+) regions and mean fluorescence intensity of COL12A1 expression was measured.

### Statistical analysis

For proteomic analysis, the Manchester Proteome Profiler suite was used which employs a range of R packages to process the protein intensity data, perform cluster analysis and identify significantly enriched proteins. A cut-off using a p-value of 0.01 (Benjamini-Hochberg method) along with a fold change of 2 was applied to determine significantly regulated proteins in each chosen pairwise comparison. The suite was further employed to perform clustering and Gene ontology (GO) enrichment analysis as well as to generate enhanced volcano and box plots to validate proteins of interest.

For immunofluorescence characterisation, invasion, TWOMBLI quantification, and Hyperion IMC COL12A1 analysis, statistical tests were performed using GraphPad Prism 10.2.2. Data were tested for normality and one-way ANOVA with Dunnett’s, Tukey’s or Holm–Sidak post hoc test used for multiple comparisons as indicated in the legends. For non-parametric data an equivalent test was used (Kruskal-Wallis with two-stage linear step-up Benjamini, Krieger, and Yekutieli procedure). Significance was determined where \*\*\**p* < 0.001, \*\**p* < 0.01 and \**p* < 0.05. For cell based assays, data represent at least three independent experiments.

## Results

### AMN and PM have an altered ECM composition compared to uninvolved regions

Tissue specimens were resected including uninvolved regions (uninvolved appendix (n=4) and uninvolved peritoneal (n=4) samples) from tumour adjacent areas and diseased regions (appendiceal mucinous neoplasms (n=3) and peritoneal metastases (n=4)). Following enrichment of insoluble matrix proteins, LC-MS/MS-based mass-spectrometry and label free quantification (Figure 1A), we identified 2171 proteins in our cohort (Supplementary Table 2). Unsupervised hierarchal clustering revealed 4 distinct clusters when comparing uninvolved appendix with AMNs and uninvolved peritoneum with PMs (Figure 1b, Supplementary Table 3). Cluster 4 identified proteins highly expressed following the development of either AMNs or PMs when compared to their respective uninvolved regions and included an increased abundance of the gel-forming mucin MUC2, and cell surface associated mucins MUC1 and MUC13. Also enriched in AMN and PM wereADAMDEC1 (a disintegrin protease), P4HB (a prolyl hydroxylase subunit essential for collagen production), tetraspanin 1 (a TSPAN protein recently associated with tumour cell invasion and migration in pancreatic cancer (30, 31)) and the proteoglycan fibromodulin (FMOD) (Figure 1C). Interestingly, the only collagen significantly differently expressed in both diseased regions compared to the uninvolved regions was a FACIT collagen, COL12A1 (Figure 1C). To identify associated pathways and functions of differentially expressed proteins in the diseased regions compared to the uninvolved, GO enrichment analysis was performed. In cluster 4, the most significantly enriched GO terms in the biological process category included organisation of both the extracellular matrix and the extracellular structure. Furthermore, in cellular component category, the most significant enrichments in cluster 4 were found in collagen-containing ECM, focal adhesions, and cell-substrate junctions. Taken together, these data suggest a key role for ECM and matrix processing in PMP.

**Figure 1:**
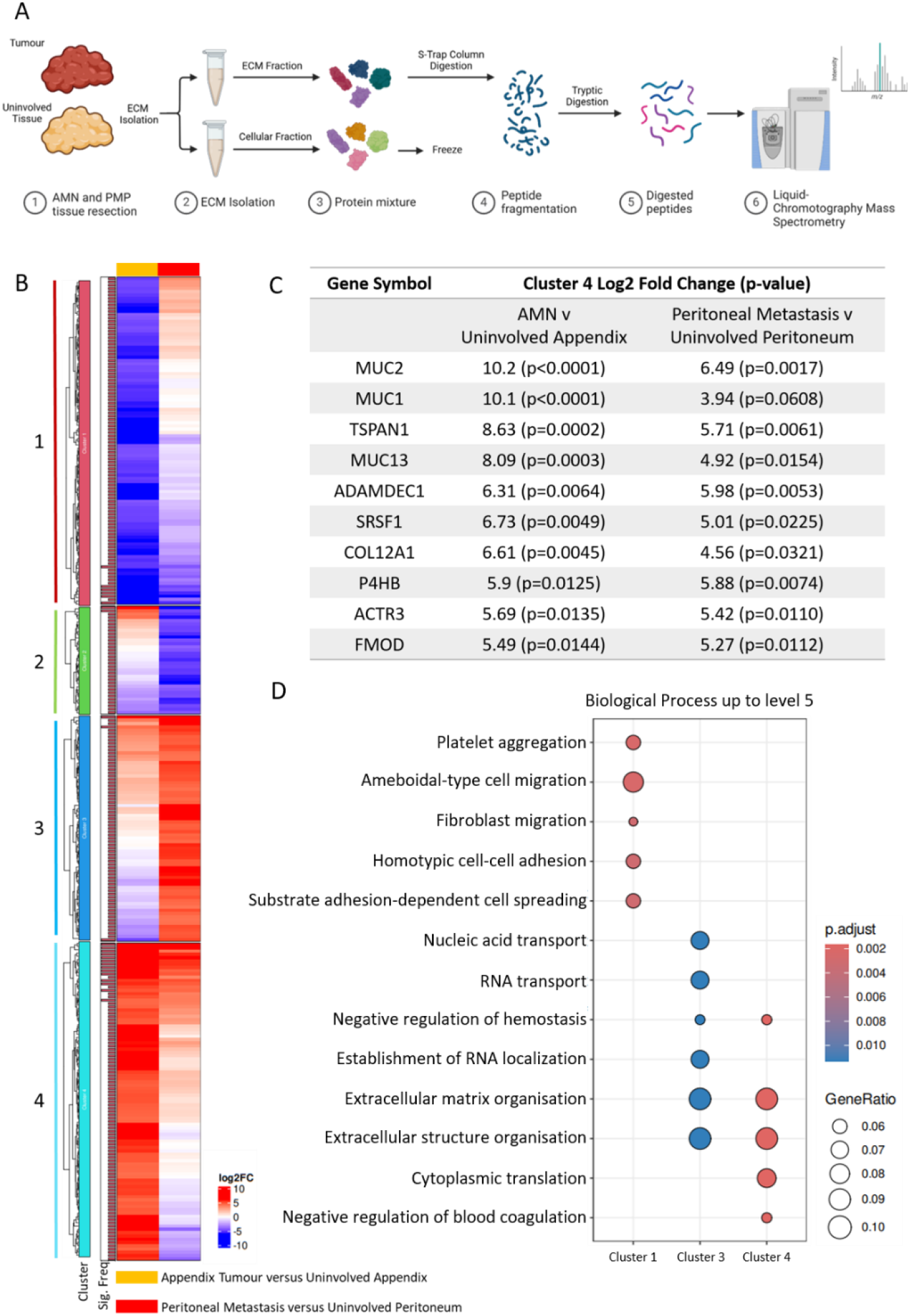
Proteomic analysis of PMP patient-derived extracellular matrix. (A) Schematic workflow for the collection of patient tissue, isolation of insoluble matrix proteins and LC-MS/MS-based proteomic analysis of the pseudomyxoma peritonei patient matrisome. (B) Cluster analysis and heatmap representation of the log2fold change of 299 differentially expressed proteins (p < 0.01). (C) Top enriched proteins in cluster 4 by at least five-fold with selected proteins labelled with gene symbols for clearness. (D) Gene ontology enrichment analysis displaying the Biological Processes between each cluster. The dot plot demonstrates enriched pathways with the colour chart represent significance and the dot circumference representing the gene ratio.

### The matrisome in AMN and AMN-PM

Given that ECM remodelling related GO pathways were identified in our global proteomic analysis (Figure 1), we explored the breakdown of matrisome proteins (core matrisome and matrisome-associated) in the diseased versus uninvolved regions. We matched all 2171 isolated proteins (Figure 1) to the matrisome to reveal a total of 315 matrisome proteins across the cohort (Supplementary Table 4). Following hierarchal clustering, 4 matrisome clusters were identified: significantly enriched in uninvolved tissues (cluster M1), significantly enriched in AMN (but not PM, cluster M2), significantly enriched in PM (but not AMN, cluster M3) and significantly enriched in both AMN and PM (cluster M4, Figure 2A).

**Figure 2:**
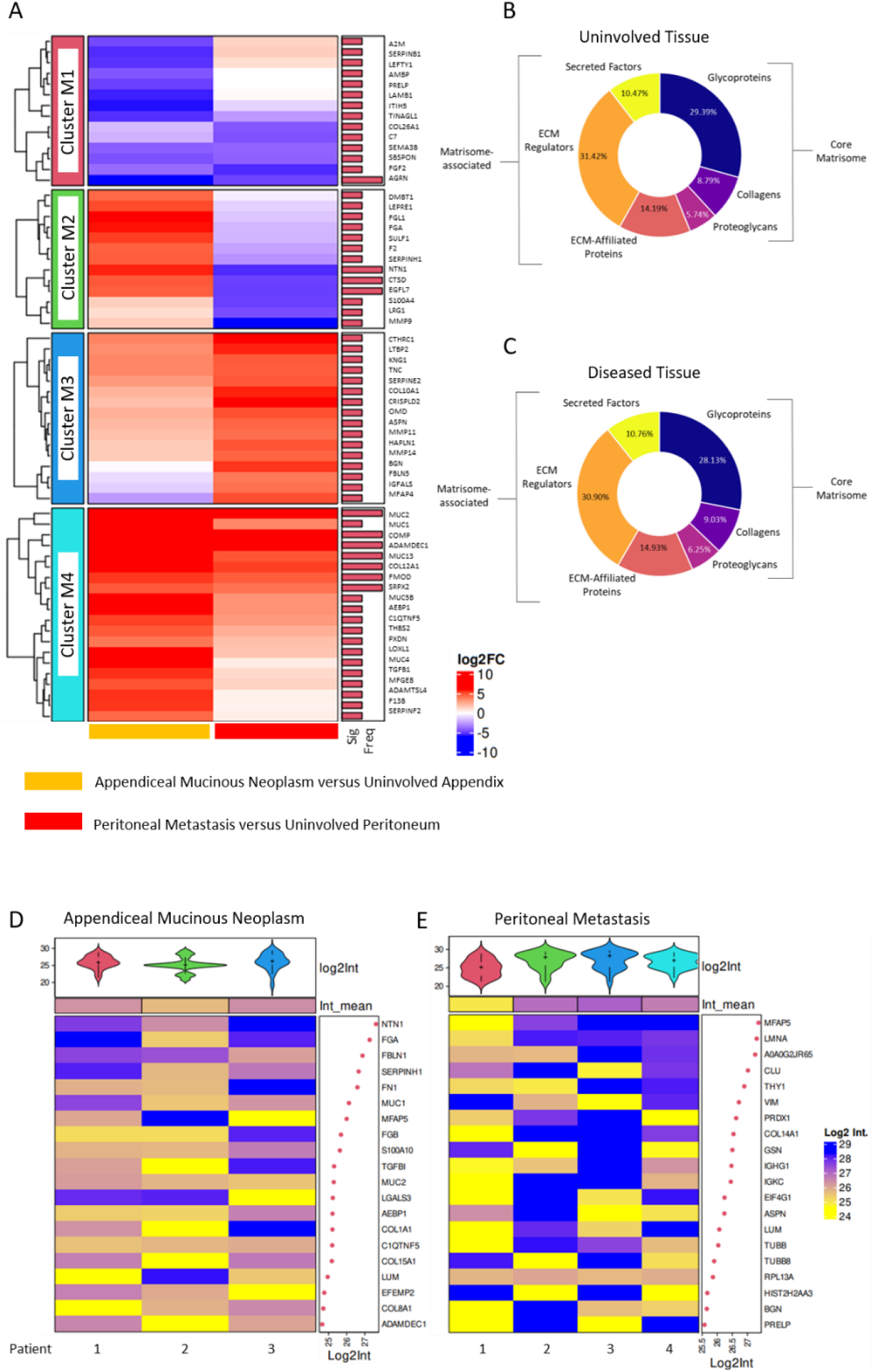
Matrisome profiling of AMN and AMN-PM. (A) Cluster analysis and heatmap representation of the log2 fold change of 63 differentially expressed proteins (p < 0.01, paired two-sided Student’s t- test with Benjamini–Hochberg correction). (B) Percentages of matrisome proteins identified in uninvolved tissues divided into matrisome subcategories. (C) Percentages of matrisome proteins identified in diseased tissues divided into matrisome subcategories. (D) Top abundant matrisome proteins in appendiceal mucinous neoplasm (E) Top abundant matrisome proteins in peritoneal metastases. Colour of abundance heatmaps represents Log2 intensity for individual patients, with dot plot representing the mean Log2 intensity across the cohort. Violin plots indicate the distribution of the log2 fold change values.

For uninvolved tissues, 130 proteins were classified as core matrisome proteins, including 26 collagens, 87 glycoproteins and 17 proteoglycans. Additionally, 166 proteins were classified as matrisome-associated proteins and included 42 ECM-affiliated proteins, 93 ECM regulators and 31 secreted proteins. For diseased tissue, 125 proteins were classified as core matrisome proteins including 26 collagens, 81 glycoproteins and 18 proteoglycans. Additionally, 163 proteins were classified as matrisome-associated proteins and included 43 ECM-affiliated proteins, 89 ECM regulators and 31 secreted proteins. This dataset represents the first detailed characterisation of matrix in AMN and PM, and whilst significant differences were identified between sample types the overall fractions of each matrisome category remained largely similar (Figure 2B, C).

To determine the dominant matrix components surrounding AMN and PMs, we identified the top 20 abundant proteins at each diseased site. In AMNs, the most abundant proteins were netrin-1 (*NTN1*), fibronectin (*Fn1*), type I collagen (*COL1A1*), AE Binding Protein 1 (*AEBP1*), and a Class II member of the small leucine-rich proteoglycan family (SLRP) - lumican (*LUM*). Furthermore, in PMs, the most abundant proteins included microfibrillar-associated protein 5 (*MFAP5*), vimentin (*VIM*), lumican (*LUM*) and another SLRP - biglycan (*BGN*).

Following the identification of abundant proteins surrounding the diseased regions, we developed 3D hydrogels that recapitulated crucial components of the tumour ECM to assess which factors influence the ability of cells to invade from tumour spheroids using the of N14A PMP cell line. When embedded into a type I collagen-based hydrogel, N14A cells rapidly invade into the surrounding matrix. The area invading cells occupied was significantly reduced upon introduction of recombinant biglycan into the collagen gels (I < 0.0001) whereas lumican had minimum influence (*p* = 0.0710) (Figure 3A, B). Interestingly, when type I collagen was combined with fibronectin, PMP cells invaded to a significantly greater extent (*p* < 0.0001), but this was attenuated when biglycan, lumican, or a combination of both SLRPs were included in the hydrogel (*p* < 0.0001). These data indicate that PMP cell invasion into interstitial matrix is heavily dependent on the specific composition of matrix proteins.

**Figure 3:**
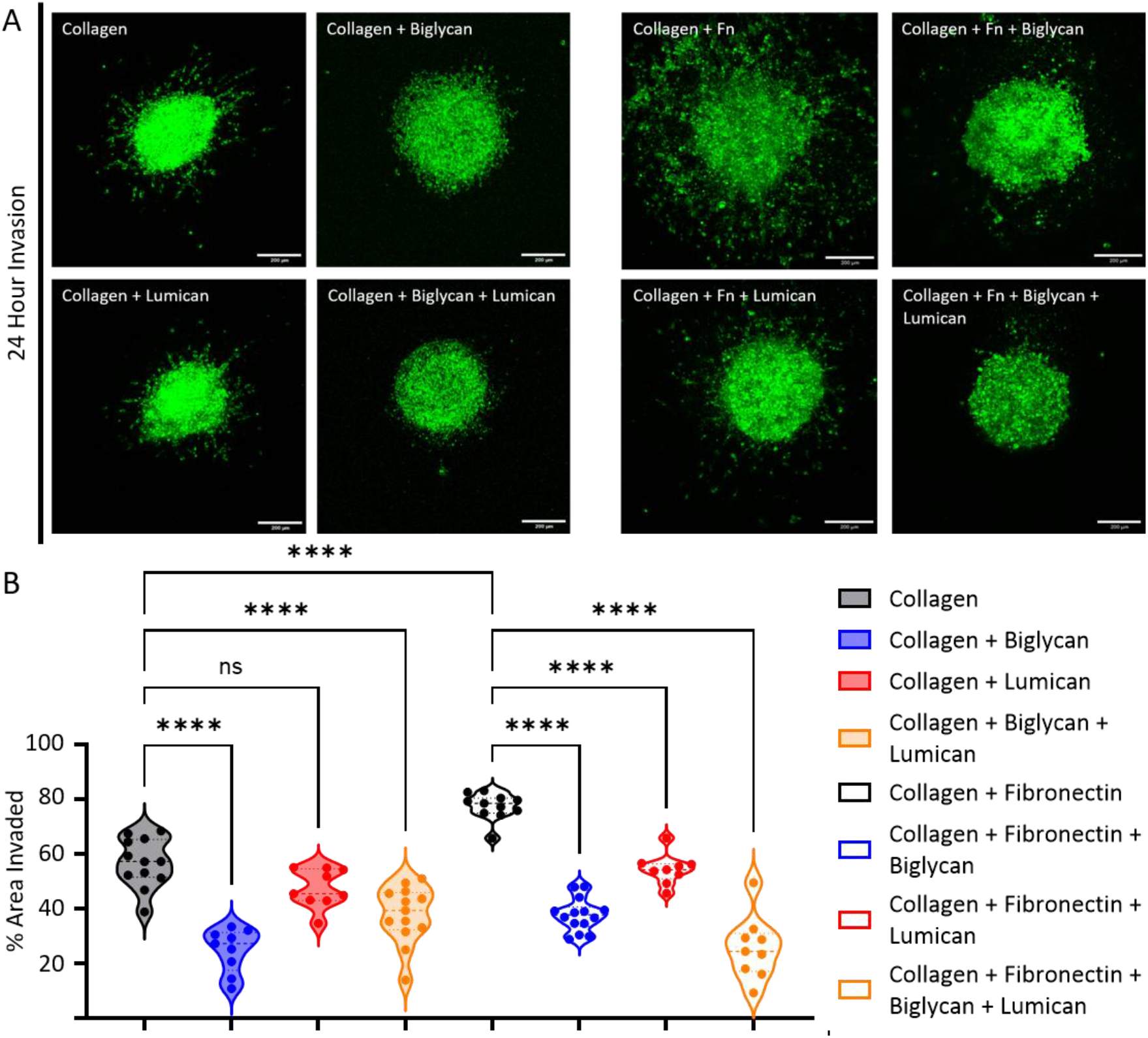
Matrix components confer pro- and anti-invasive properties to the AMN and PMP niche. (A) N14A spheroid invasion assay showing N14A spheroids seeded into collagen gels supplemented with combinations of biglycan, lumican and fibronectin (FN) for 24 hours and treated with EGF/HGF. (B) Quantification of percentage invasion area of N14A spheroids at 24 hours in relation to core spheroid area. One-way ANOVA with Holm–Sidak post hoc test was used for multiple comparisons. Significance was determined where **** denotes p < 0.0001, *** denotes p < 0.001, ** denotes p < 0.01, * denotes p < 0.05 and ns denotes p > 0.05 (not significant). Data represent at three independent experiments. Scale bar represents 200 μm.

### Collagen remodelling in the AMN and PM niche

The ECM provides a structural scaffold for tissues, with fibrillar collagens at the core of many of these support mechanisms. However, a dynamic and remodelled ECM is critical for the development and progression of various cancer types resulting in unfavourable phenotypic characteristics and a reduction in treatment responses. Quantifying and understanding the architecture of these fibrillar-based matrices surrounding and supporting PMP could lead to novel insight into how primary AMNs disseminate from the appendix and develop metastatic niches within the peritoneal cavity. To investigate this, we stained formalin-fixed paraffin embedded (FFPE) tissue from patients with PMP, including both uninvolved and diseased regions, with picrosirius red and subjected the tissue to polarised light to visualise collagens due to their birefringent properties (Figure 4A). Quantitative analysis was performed using the TWOMBLI plugin within ImageJ (29) to evaluate collagen fibre density and organisation (Figure 4B, Supplementary Figure 2). Peritoneal metastases differed significantly from other tissues studied across several metrics, and AMN tissues showed significantly increased alignment suggesting that collagen remodelling and reorganisation is a feature of AMN and resulting peritoneal metastasis (Figure 4A). Within the peritoneal metastases, increased deposition of fibrillar collagen was observed when compared to the primary AMNs and the uninvolved peritoneum (collagen area, Figure 4A, B). Furthermore, peritoneal metastases were composed of an increased percentage of high-density matrix when compared to the primary AMNs and had a lower lacunarity (a measure of how collagen fibres fill the space, with lower scores indicating a reduction in matrix gaps, Figure 4A, B). When assessing collagen organisation within the peritoneal metastases, we observed an increase in hyphal growth unit (number of collagen fibre endpoints per fibre length) when compared to uninvolved peritoneum or appendix and AMNs. Additionally, peritoneal metastases exhibited a more similar (fractal dimension) and well aligned collagen matrix when compared to the uninvolved peritoneum and AMNs (Figure 4A, B). These data demonstrate that collagen deposition and remodelling is a significant feature of AMN and is even more apparent in PMs, suggesting that this could contribute and support disease progression.

**Figure 4:**
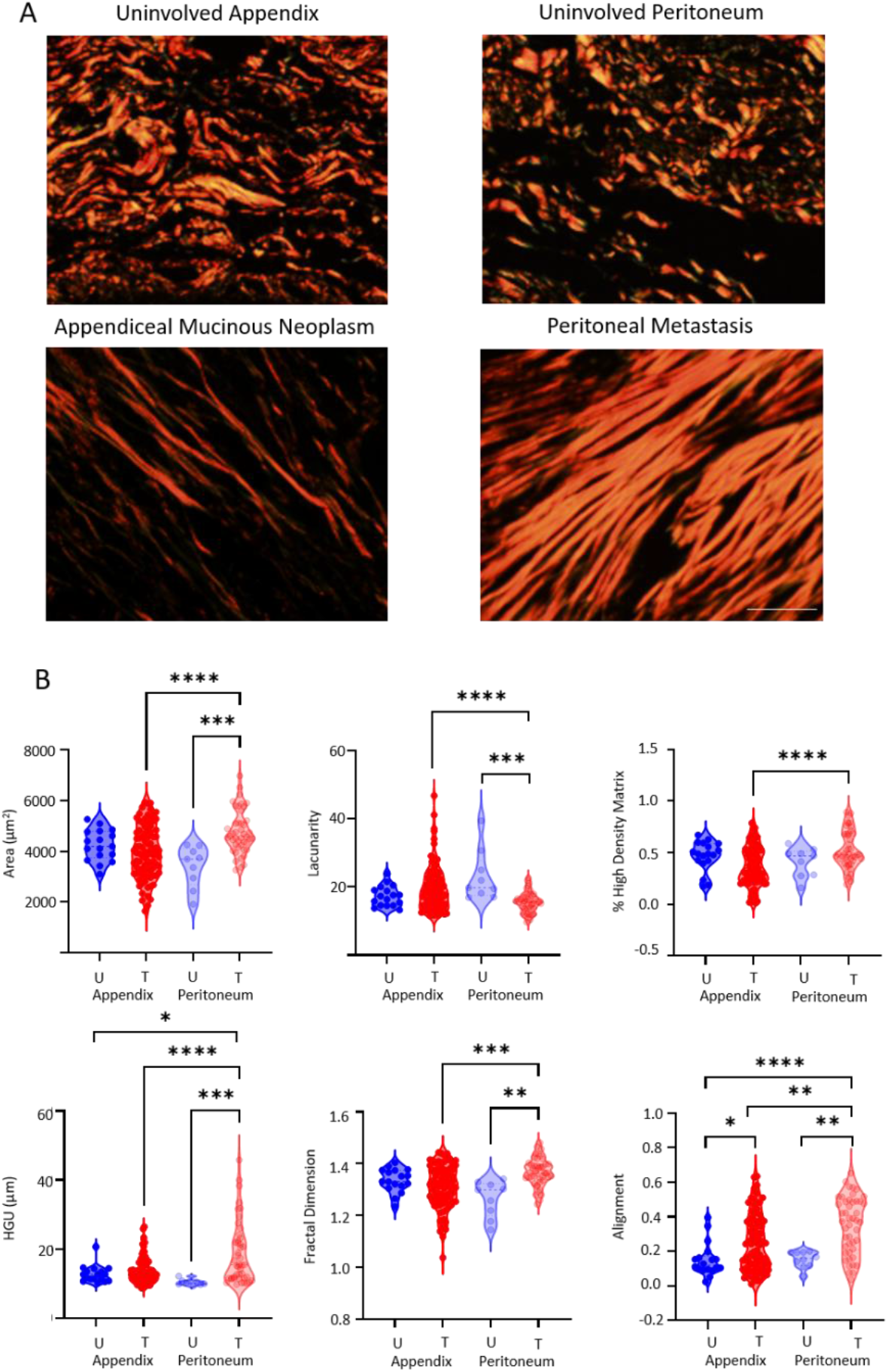
Peritoneal metastases arising from appendiceal mucinous neoplasms demonstrate a dynamic and remodelled extracellular matrix. (A) Representative picrosirius red staining on FFPE sections of uninvolved appendix, uninvolved peritoneum, appendiceal mucinous neoplasm, and peritoneal metastases subjected to polarised light. (B) Violin plots denoting quantification of TWOMBLI matrix patterns including area, lacunarity, percentage of high-density matrix, hyphal growth unit (HGU), fractal dimension and alignment. U = uninvolved and T = tumour. One-way ANOVA with Tukey’s post hoc test was used for multiple comparisons or an equivalent non-parametric test for Lacunarity, HGU and Alignment. Significance was determined where **** denotes p < 0.0001, *** denotes p < 0.001, ** denotes p < 0.01, * denotes p < 0.05 and ns denotes p > 0.05 (not significant). Each data point depicts an independent reading, taken in triplicate per ROI. Three ROIs per tissue section were selected based on their locality to the tumour-stroma boundary per patient (3 for uninvolved appendix/ appendiceal mucinous neoplasm/ peritoneal metastases; 1 for uninvolved peritoneum). Scale bar represents 200 μm.

### Collagen XII (COL12A1) is upregulated in AMNs and their peritoneal metastases

Stromal factors that could drive AMN, and adaptation of the metastatic niche within the peritoneal cavity during disease progression are poorly understood. We therefore sought to determine the major differences in matrix composition between diseased versus uninvolved tissue. We compared expression changes of matrisome proteins in the diseased regions when compared to the uninvolved regions (Figure 5). Within the appendiceal mucinous neoplasms, highly significant upregulation of a number of both cell-surface and secreted mucins were observed compared to the uninvolved appendix (*MUC2, MUC1, MUC13, MUC5B*) along with a number of core matrix glycoproteins (*COMP, AEBP1, SRPX2, DMBT1*) (Figure 5A, C). Interestingly, there was also a significant loss of basement membrane proteins (*ARGN, LAMB1, PRELP, TINAGL1*) suggesting a breakdown of the basement membrane during the development of AMNs within the appendix (Figure 5A, C). Within the peritoneum, we observed a significant upregulation of numerous proteins involved in tumour progression and metastasis (*CTHRC1, LTBP2, HAPLN1, ASPN, COL10A1*) along with regulators and modulators of the ECM within the TME (*COMP, CRISPLD2, ADAMDEC1, BGN, MMP2*) (Figure 5B). Interestingly, collagen XII (COL12A1) was the only collagen within our cohort that exhibited significant upregulation in both the primary AMNs (Log_2_ fold change=4.56) and peritoneal metastases (Log_2_ fold change=6.61) when compared to the uninvolved regions (Figure 5).

**Figure 5:**
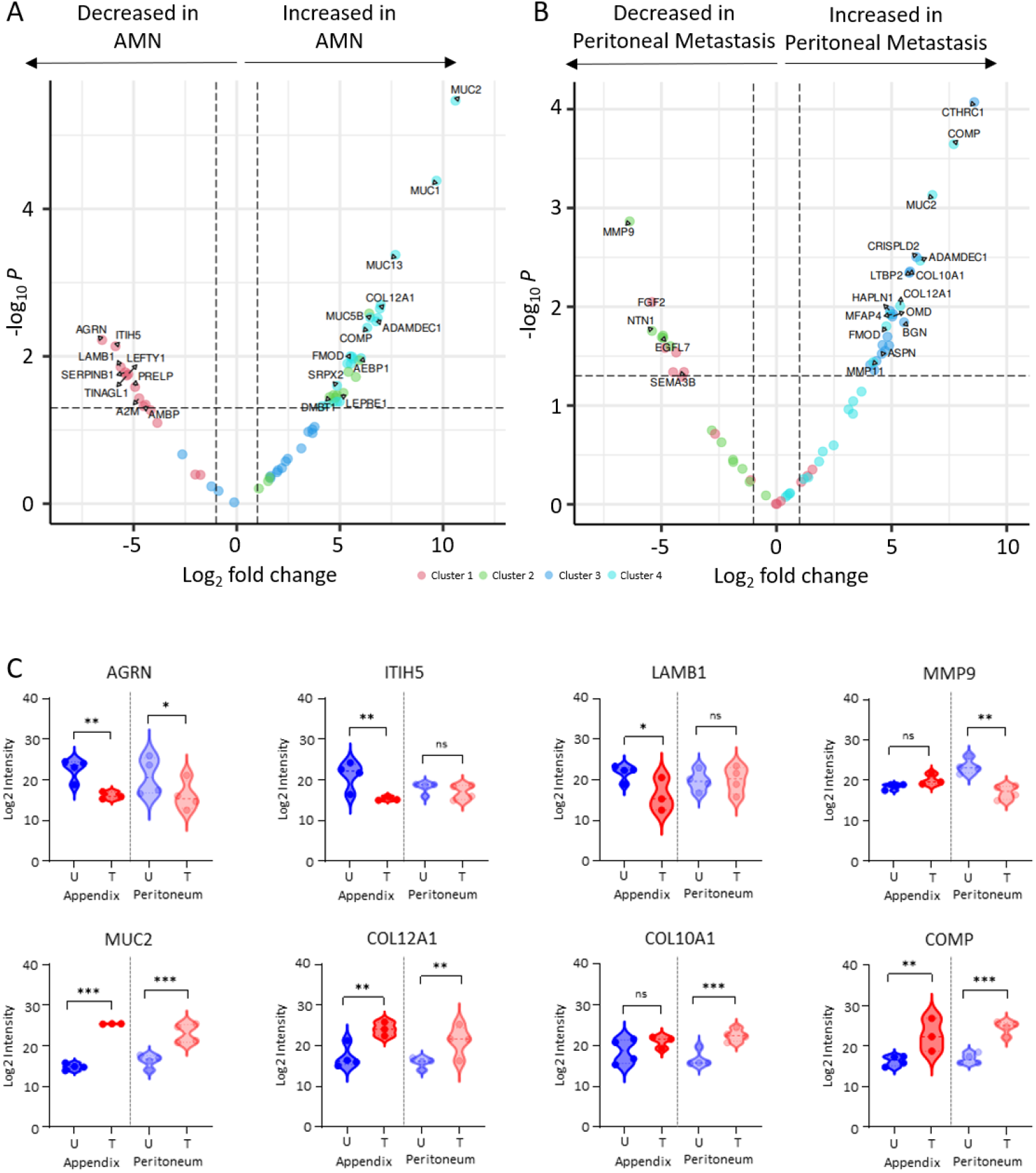
COL12A1 is significantly enriched within both the primary appendiceal mucinous neoplasms (AMN) and peritoneal metastases when compared to uninvolved appendix and peritoneum. (A) Volcano plot of appendix tissue matrisome proteins quantified by LC-MS/MS proteomics. Significantly differentially regulated proteins enriched or depleted in AMN by at least two-fold are to the side of the dashed cut-off line. (B) Volcano plot of peritoneal tissue matrisome proteins quantified by LC-MS/MS proteomics. Significantly differentially regulated proteins enriched or depleted in peritoneal metastases by at least two-fold are to the side of the dashed cut-off line. Coloured data points represent the clusters within the cohort and selected proteins are labelled with gene names for clearness. (C) Violin plots of selected significantly regulated proteins showing log2 intensities. Unpaired T-tests were used to determine significant differences between uninvolved (U) versus tumour (T) appendix and U versus T peritoneum. Significance was determined where **** denotes p < 0.0001, *** denotes p < 0.001, ** denotes p < 0.01, * denotes p < 0.05 and ns denotes p > 0.05 (not significant).

Using multiplexed imaging mass cytometry, we determined the expression of COL12A1 within PMP patients (Figure 6A) (Supplementary Table 5). Using smooth muscle actin (SMA) as a marker of the stromal compartment and CK20 as a marker of tumour and/or epithelial cells within the appendix and peritoneum, we observed a significantly higher mean fluorescence intensity of COL12A1 within the stromal compartment of AMNs and PMs when compared to their respective uninvolved regions (Figure 6B). COL12A1 expression was relatively high in epithelial cells of the appendix (Figure 6C), and in fact levels were decreased in epithelia of AMN patients, whereas the peritoneum is largely devoid of epithelia until onset of PMP. Taken together, these data suggest that there is a significant increase in COL12A1, specifically within the stromal compartment in AMN and PM, and that this could contribute to the ability of AMNs to progress.

**Figure 6:**
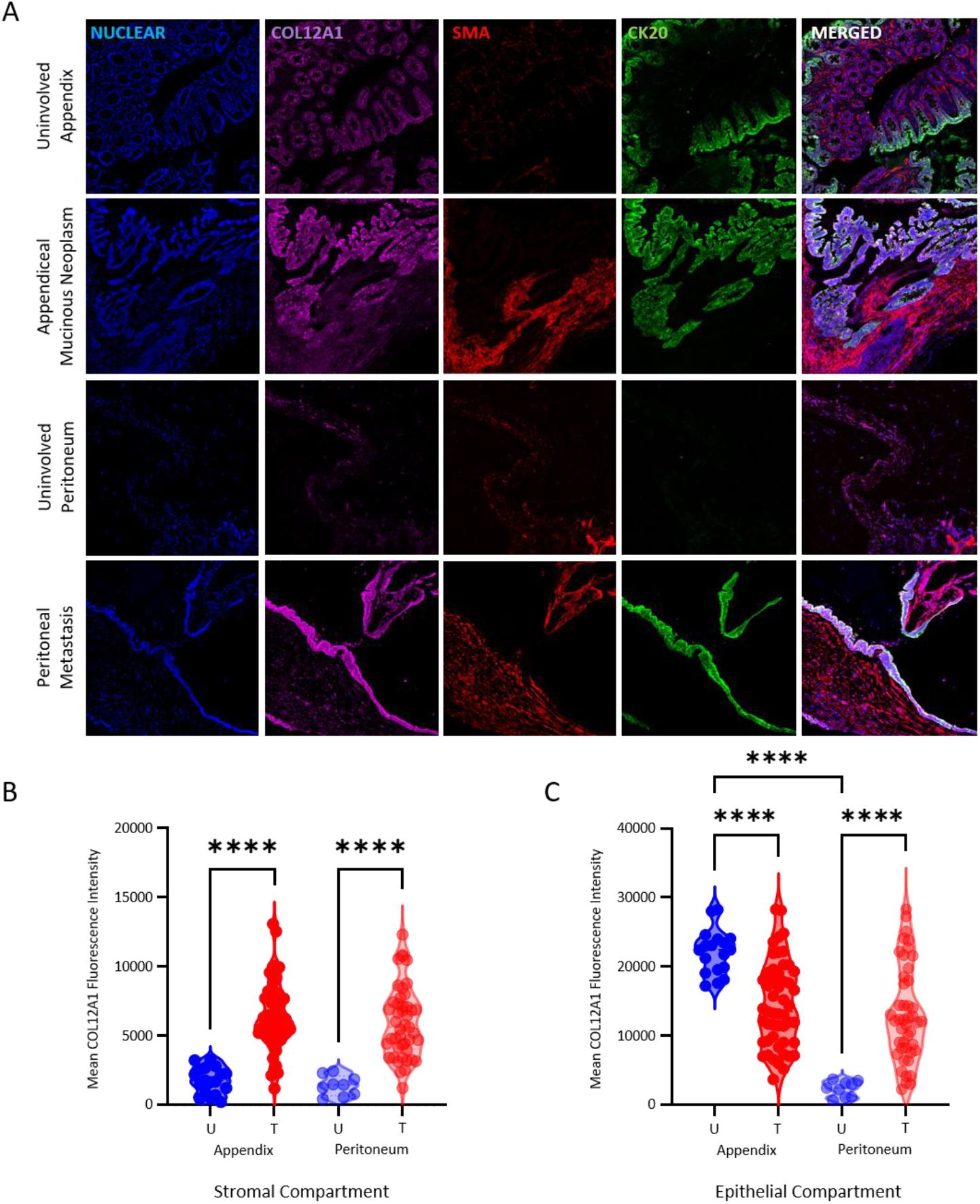
COL12A1 expression increases within stromal regions of AMN and PMs increases. (A) Multiplexed images demonstrating COL12A1 expression in both the stromal (SMA+) and epithelial (CK20+) compartments of uninvolved appendix, appendiceal mucinous neoplasms, uninvolved peritoneum and peritoneal metastases. (B) Violin plots indicating significantly higher expression of COL12A1 within the stromal compartments of appendiceal mucinous neoplasms and peritoneal metastases when compared to their respective uninvolved regions. (C) Violin plots indicating significant differences in COL12A1 expression within the epithelial compartments of uninvolved versus diseased regions. U = uninvolved and T = tumour. One-way ANOVA with Holm–Sidak post hoc test was used for multiple comparisons. Significance was determined where **** denotes p < 0.0001, *** denotes p < 0.001, ** denotes p < 0.01, * denotes p < 0.05 and ns denotes p > 0.05 (not significant). Each data point depicts three representative areas selected based on their locality to the tumour-stroma boundary across a minimum of three regions of interest per patient. Patients n = 2 uninvolved appendix, n = 5 appendiceal mucinous neoplasm, n = 1 uninvolved peritoneum, and n = 5 peritoneal metastasis. Scale bar represents 200 μm.

### COL12A1 expression in CAFs associates with increased PMP cell invasion

COL12A1 is known for its ability to regulate the organisation and biomechanics of type I collagen and has been associated with collagen I bundling and matrix stiffness within aggressive breast cancer (32). More recently, through single-cell transcriptomics, COL12A1 has been shown to be secreted by CAFs resulting in type I collagen reorganisation and the development of a pro-invasive TME supporting breast cancer metastatic dissemination (17). As COL12A1 within our cohort has a higher expression within the stromal compartments of diseased regions when compared to the uninvolved regions, we isolated PM-derived CAFs from patients with histologically confirmed PMP. CAF myofibroblast-like phenotype was confirmed using SMA and FAP (Supplementary Figure 3A and B), and COL12A1 expression was observed within these cell cultures (Supplementary Figure 3C and D). When co-cultured with COL12A1 positive CAFs from 2 different patients, we observed significant increases in N14A PMP cell invasion into collagen-based hydrogels when compared to N14A monocultures alone, suggesting that CAFs facilitate PMP cell invasion. COL12A1 expression was then silenced within the 2 CAF lines using two independent siRNAs (siCOL12A1_1 and siCOL12A1_2), and the ability of CAFs to support PMP cell invasion quantified. COL12A1 silenced CAFs from the 2 patients showed a significant decrease invasion of PMP cells, suggesting that COL12A1 expression in stromal CAFs remodels collagen I to enhance the invasive migration of PMP cells (Figure 7A-D).

**Figure 7:**
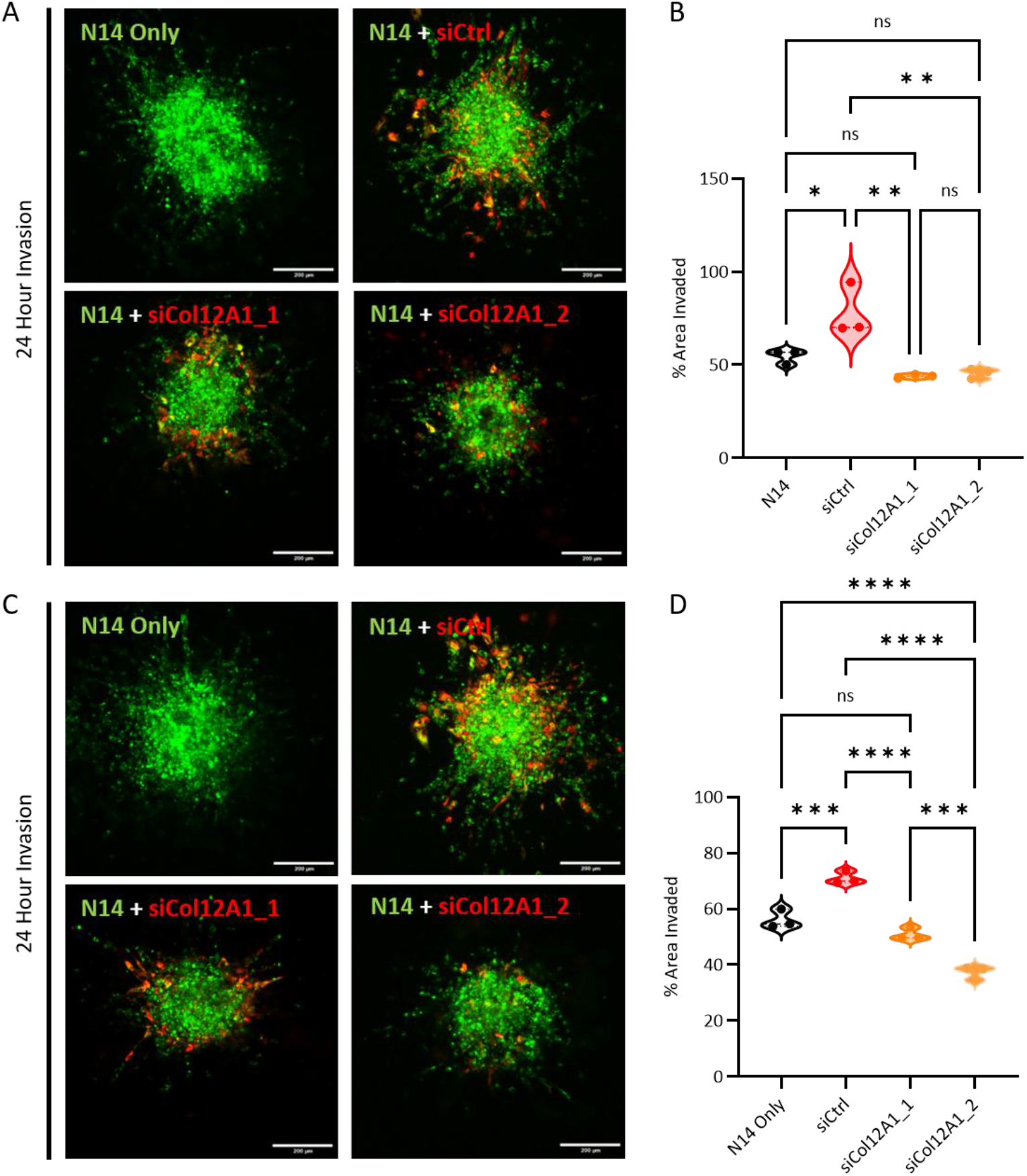
COL12A1 expression in patient-derived fibroblasts facilities PMP cell invasion in3D spheroid models. (A) N14A cells co-cultured with patient 1-derived fibroblasts spheroid invasion assay showing non-silenced COL12A1 (siCtrl) fibroblasts increased N14A cell invasion whereas siCol12A1_1 and siCol12A1_2 knockdown in fibroblasts reduced its ability to promote N14A invasion over 24 hours. (B) Violin plots showing quantification of the effects patient 1-derived fibroblasts on N14A invasion in 3D spheroid models. (C) N14A cells co-cultured with patient 2-derived fibroblasts spheroid invasion assay showing siCtrl fibroblasts increased N14A cell invasion whereas siCol12A1_1 and siCol12A1_2 knockdown in fibroblasts reduced its ability to promote N14A invasion over 24 hours. (D) Violin plots showing quantification of the effects patient 2-derived fibroblasts on N14A invasion in 3D spheroid models. One-way ANOVA with Tukey’s post hoc test was used for multiple comparisons. Significance was determined where **** denotes p < 0.0001, *** denotes p P< 0.001, ** denotes p < 0.01, * denotes p < 0.05 and ns denotes p>0.05 (not significant). Each data point depicts the mean of an independent experiment n = 3. Scale bar represents 200 μm.

## Discussion

A dynamic and remodelled ECM is a key driver of many solid tumours and at the forefront of metastatic niche development within the peritoneal cavity (33). PMP is thought to be driven by KRAS and GNAS mutations, along with changes to the tumour microenvironment, with secretion of mucins MUC2A and MUC5AC being a defining feature of this disease (34). However, tumour extrinsic stromal factors are less associated with disease progression. Recently, CAF transcriptional signatures have been found to increase the predictive power of tumour transcriptional profiling (35), but associated changes in insoluble matrix are under-investigated. Our interrogation of the matrix landscape of PMP has determined that matrix composition in the TME helps to shape the invasive behaviour of tumour cells. We demonstrate that stromal COL12A1 upregulation is a feature of AMN and PM, with CAF-generated COL12A1 facilitating invasive migration of PMP cells. Furthermore, we uncovered critical morphological properties within AMN and PM stromal architecture, suggesting both matrix composition and its organisation are implicated within peritoneal metastatic dissemination.

To date, few models have been developed that accurately recapitulate the 3D architecture, stromal interactions, and underlying biology needed to investigate mechanisms and therapeutic targets. We therefore set out to investigate the AMN and PM niche using a proteomic approach, and initial gene ontology analysis indicated that extracellular matrix organisation is a key feature of the protein networks upregulated in AMN and PM (Figure 1D). Further analysis of the matrisome highlighted an abundant, collagen-rich matrix associated with AMN and PM and their equivalent uninvolved tissues. Amongst the most abundant proteins identified were COL1A1, the major component of fibrillar collagen-I matrix, as well as the glycoprotein FN and SLRPs lumican and biglycan. Interestingly, our results indicate that COL1A1 and FN (found abundantly in AMN tissue) might be pro-invasive, whereas biglycan (high levels in PMs, and to an extent lumican) acts as to suppress invasive migration of N14A PMP cells. We suggest that the matrix niche surrounding AMN is conducive to escape of cells (for example following appendix rupture), whereas factors (e.g. biglycan) in the peritoneal niche could help to generate a barrier that prevents further spread.

Analysis of the matrisome altered between uninvolved and diseased tissues indicated that the basement membrane, a lattice-like sheet of matrix that serves to separate tissue compartments, was downregulated in AMNs (Figure 2A, cluster M1) and may therefore be reduced early in disease progression. Key components of the basement membrane were down regulated including LAMB1 (laminin β1, a major component of the basement membrane lattice), PRELP (Proline and Arginine Rich End Leucine Rich Repeat Protein) and AGRN (Agrin), which function to anchor the basement membrane to connective tissue and contribute to the structural integrity and stability of the basement membrane (36, 37). Reduced basement membrane integrity would be expected to weaken the barrier between AMNs and their surrounding environment, and when couple with an abundant collagen and fibronectin-rich stromal matrix could help facilitate dissemination. Expression of PRELP is significantly downregulated in colorectal cancer (CRC) with its expression negatively correlating with lymph node metastases and poor prognosis (38). Furthermore, destabilisation of ECM through loss of ITIH5 (Inter-alpha-trypsin inhibitor heavy chain 5) and inhibition of integrin/FAK signalling pathway via TINAGL1 downregulation may represent a specific matrix remodelling pathway associated with early transformation of the basement membrane and dissemination into the peritoneal cavity (39). Proteins upregulated in AMN (Figure 2A cluster M2, Figure 5A)) included DMBT1 (Deleted in Malignant Brain Tumours 1, a secreted mucin-like glycoprotein (40)), LEPRE1 and SERPINH1 (involved in collagen biosynthesis (41)) and other matrix modifying enzymes (ADAMDEC1, CTSD (42)). Within the PM samples, cluster M3 revealed upregulated proteins when compared to the uninvolved peritoneum. High abundance of CTHRC1 (Collagen Triple Helix Repeat Containing 1) was observed, a secreted glycoprotein often upregulated in solid tumours such as CRC, with CTHRC1 correlating with hepatic metastases and macrophage infiltration through TGF-β signalling (43). COL10A1 abundance was also upregulated within PMs, a secreted collagen which promotes gastric cancer cell epithelial-mesenchymal transition (EMT) and invasion via TGF-β1-SOX9 axis (44). This is of particular interest, suggesting COL10A1 expression may specifically be associated with the remodelling of the peritoneal niche seen in PMP.

Several matrix proteins were highly abundant proteins within both AMNs and PMs (cluster M4), with elevated secreted MUC2 observed. This gel-forming mucin is linked with advanced disease and poor prognosis in many cancer types, associating with peritoneal chemoresistance and immune infiltration (45). In addition, COL12A1 was the only collagen significantly upregulated within both AMNs and PMs. Collage XII is a FACIT collagen involved in regulating the spatial organisation and bundling of type I collagen fibrils by connecting them to the ECM. It has been previously shown that CAFs are key secretors of collagen XII, and its expression correlates with high mammographic density and poor clinical outcomes for women with breast cancer (32). Interestingly, COL12A1 expression has been implicated in immune surveillance through its effects on collagen I architecture (46). Elevated COL12A1 expression in breast cancer predicts poor response to anti-PD-1/PD-L1 therapy, correlating with M2 macrophage infiltration and reduced overall survival (47). Furthermore, CAF-secreted collagen XII has shown to promote breast cancer metastasis to the lung and support an invasive TME through collagen I remodelling (17). Many solid tumours have highlighted CAFs as key manipulators of the metastatic niche, driving fibrosis and the increased deposition of a malignant stroma. The recruitment of fibroblasts into the peritoneal cavity is promoted by integrin α5 expression on high-grade serous ovarian cancer cells within the peritoneal fluid (48). Furthermore, these peritoneal metastasis-associated fibroblasts express elevated levels of collagen and matrix metalloprotease 2 (MMP2) when compared to fibroblasts in adjacent uninvolved peritoneum. Following image analysis within the stroma compartments, we reveal strong expression of COL12A1 in AMNs and PMs, with CAFs isolated from PMs expressing COL12A1. Importantly, co-culturing of PMP cell lines with COL12A1 expressing CAFs promoted invasion within a 3D collagen-based hydrogel when compared to PMP monocultures, which was attenuated following COL12A1 knockdown. The presence of COL12A1 both at the primary and metastatic sites suggests that COL12A1 acts at different stages/ sites of the disease, promoting an invasive tumour microenvironment and potential mechanistic model underlying the dynamic ECM seen in PMP.

Along with compositional alterations of the matrix, variation in matrix architecture during tumourigenesis is widely implicated in promoting cancer progression, metastatic dissemination, and reduced treatment response. Collagen fibril organisation, structure, and abundance promotes a cancerous ECM through progressive linearization and crosslinking to stiffen the matrix and form networks for cell migration and signalling (49). A stiffened ECM changes the mechanical properties and drives disease progression in various solid tumours including pancreatic ductal adenocarcinoma (50). Our analysis of collagen organisation indicates that the fibrillar collagen matrix becomes more aligned in AMN and PMs, consistent with findings in other cancers. These data also suggest that the matrix environment surrounding PMs become more dense and homogenous, similar to characteristics seen in aggressive triple negative breast cancer (51). A more aligned collagen matrix at the invasive front of many primary tumours correlates with poor prognosis and can provide a structural pathway for cancer cell invasion (52, 53). Since collagen XII regulates collagen I fibril packing and assembly, we propose that COL12A1-expressing stromal fibroblasts, either at the primary tumour or at the peritoneal site, can remodel collagen architecture and potentially create a more permissive niche for tumour invasion and colonisation.

In conclusion, we reveal dynamic ECM changes both within AMNs and PMs, presenting potential therapeutic stromal targets at both early and late-stage disease. The elevated expression of COL12A1 at critical stages of disease progression, its potential role in early dissemination of cells to the peritoneum, and its significant association with PMP invasion *in vitro*, hails the exploration of COL12A1 as a future mechanistic lead for patients with PMP.

## Acknowledgments

PMPnet group

The following authors are participants (listed alphabetically by country and place) of the PMPnet Group: Italy: Chiorino Giovanna, Mazzarotto Francesco, Mello Grand Maurizia, Ostano Paola, Peraldo Neia Caterina, Fondazione Edo ed Elvo Tempia, Biella; Biletta Elena, D’Andrea Mariangela, Pisacane Alberto, Ospedale degli Infermi, Biella; Cascardi Eliano, Vaira Marco, Istituto di Candiolo-Fondazione del Piemonte per l’Oncologia-IRCCS, Candiolo; Cattaneo Laura, Ciniselli Chiara, Colletti Gaia, Illescas Oscar, Pizzamiglio Sara, Sabella Giovanna, Verderio Paolo, Fondazione IRCCS Istituto Nazionale dei Tumori, Milan; Cantarella Daniela, Isella Claudio, Medico Enzo, Università degli Studi di Torino, Turin. Spain: Bijelic Lana, Palacio Lina, Sabia Domenico, Hospital de Sant Joan Despí Moisès Broggi, Barcelona; Cabot Romero Débora, Palmer Hector G, Vall d’Hebron Institute of Oncology, Barcelona. United Kingdom: Aziz Omer, Barriuso Jorge, Chakrabarty Bipasha, Garva Richa, O’Dwyer Sarah, The Christie NHS Foundation Trust, Manchester; Barber Jill, Demonacos Constantinos, Fry Glenys, Hasan Awen, McAllister Milly, Nagaraju Raghavendar, Rogan Jane, Rostami Amin, Vasilogianni Areti, University of Manchester, Manchester.

---

**Supplementary Figure 1:**
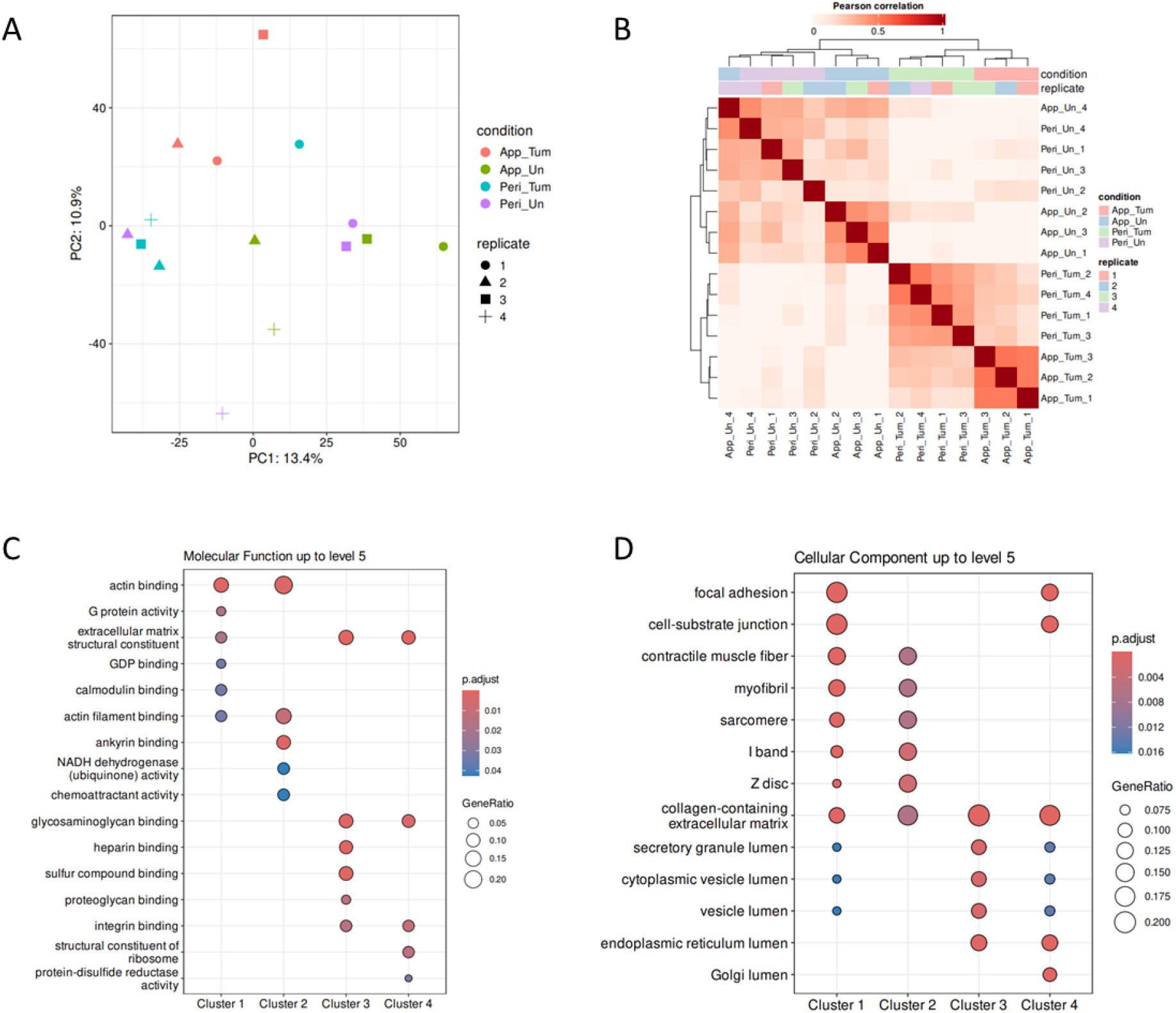
Proteomics experiment summary. (A) Principal component analysis (PCA) plot demonstrating the relationship between replicates of appendiceal mucinous neoplasm (App_Tum), uninvolved appendix (App_Un), peritoneal metastases (Peri_Tum) and uninvolved peritoneum (Peri_Un). (B) Pearson correlation matrix showing the linear relationship between conditions and replicates. (C) Gene ontology enrichment analysis displaying the Molecular Functions between each cluster. (D) Gene ontology enrichment analysis displaying the Cellular Components between each cluster. The dot plot demonstrates enriched pathways with the colour chart represent significance and the dot circumference representing the gene ratio.

**Supplementary Figure 2:**
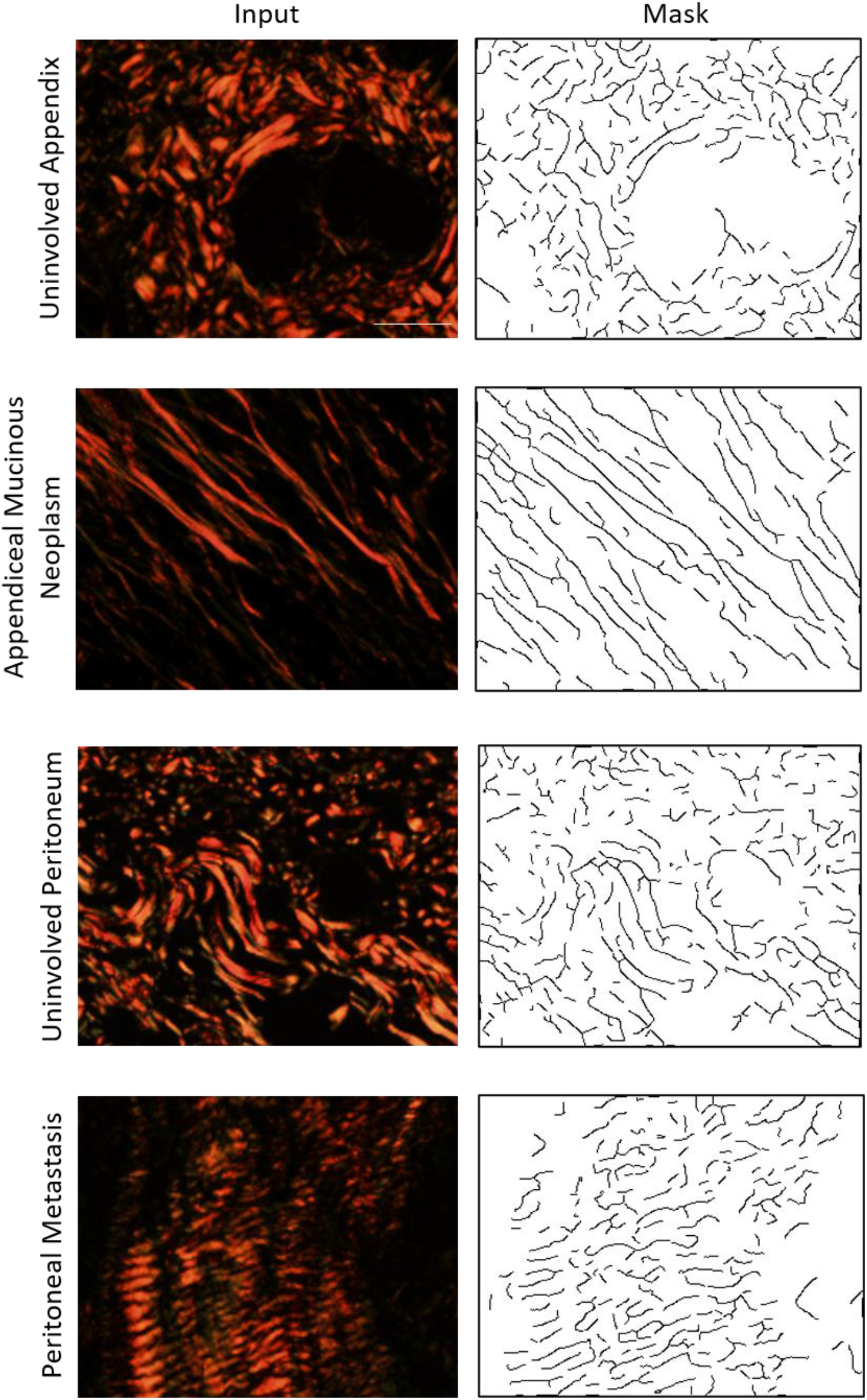
Input and outputs of TWOMBLI pipeline. Example of input images from picrosirius red stained FFPE tissues from patients with PMP, that have been subjected to polarised light, including uninvolved appendix, appendiceal mucinous neoplasm, uninvolved peritoneum and peritoneal metastasis. Output from TWOMBLI pipeline consists of masks containing matrix metrics. Scale bar represents 200 μm.

**Supplementary Figure 3:**
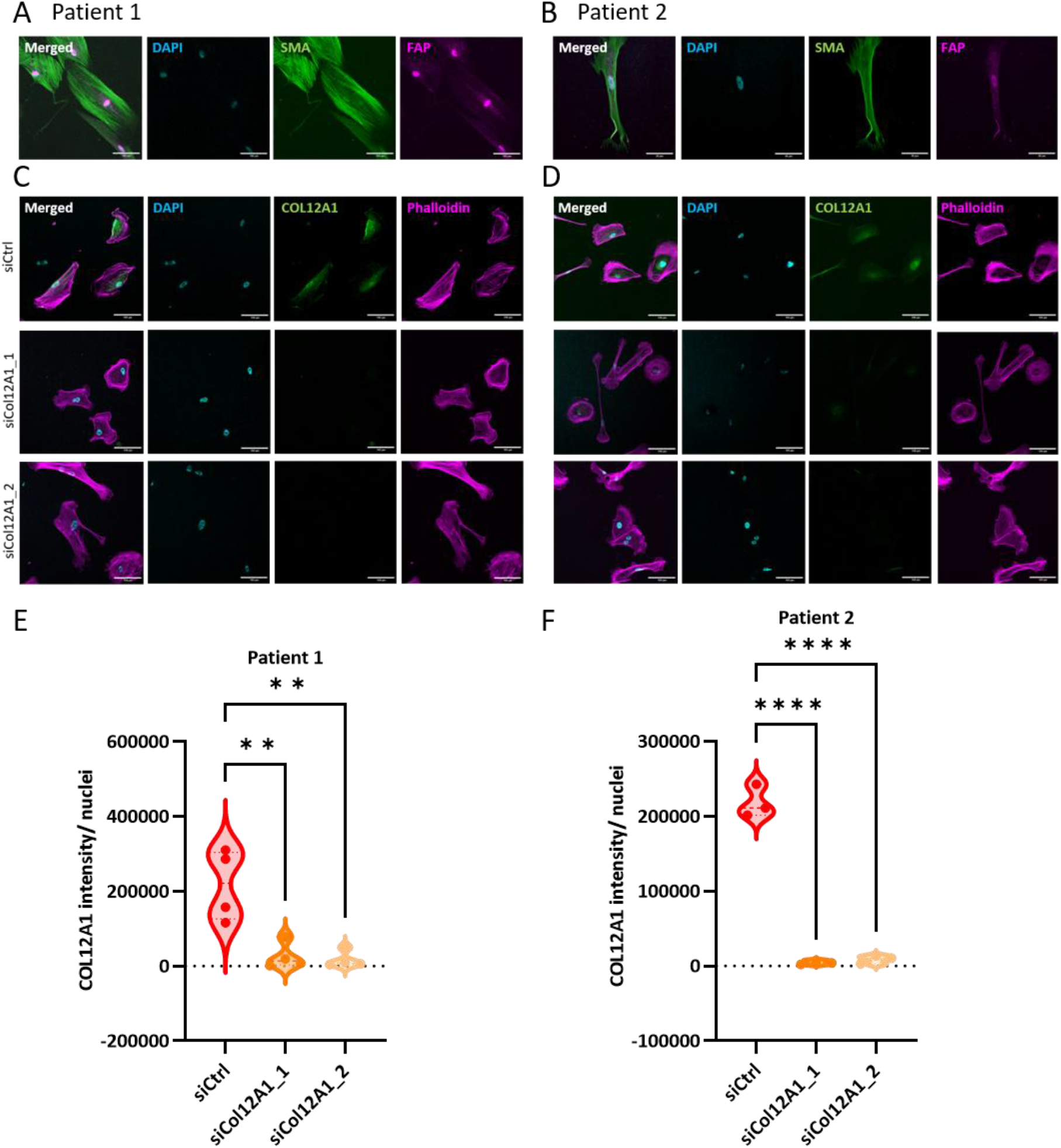
COL12A1 expression was significantly knockdown in patient 1- and patient 2-derived fibroblast cultures following SiRNA transduction. (A and B) Immunofluorescence characterisation of α-smooth muscle actin and fibroblast activation protein following transduction with siCtrl, siCol12A1_1and siCol12A1_2 in patient 1 (A) and patient 2 (B). (C and D) Immunofluorescence characterisation of COL12A1and phalloidin following transduction with siCtrl, siCol12A1_1and siCol12A1_2 in patient 1 (C) and patient 2 (D). (E and F) Violin plots displaying quantification of COL12A1 intensity relative to the number of nuclei per field of view. One-way ANOVA with Dunnett’s test was used for multiple comparisons. Significance was determined where **** denotes p < 0.0001, *** denotes p < 0.001, ** denotes p < 0.01, * denotes p < 0.05 and ns denotes p > 0.05 (not significant). Scale bar represents 100 μm.

